# A technique for resolution assessment in blind-SIM experiments

**DOI:** 10.1101/2021.09.01.458550

**Authors:** Imen Boujmil, Emmanouil Xypakis, Giancarlo Ruocco, Marco Leonetti

## Abstract

Super resolution techniques are an excellent alternative to wide field microscopy, providing high resolution also in (typically fragile) biological sample. Among the various super resolution techniques, Structured Illumination Microscopy (SIM) improve resolution by employing multiple illumination patterns to be deconvolved with a dedicated software. In the case of blind SIM techniques, unknown patterns, such as speckles, are used, thus providing super resolved images, nearly unaffected by aberrations with a simplified experimental setup. Scattering Assisted Imaging, a special blind SIM technique, exploits an illumination PSF (speckle grains size), smaller than the collection PSF (defined by the collection objectives), to surpass the typical SIM resolution enhancement. However, if SAI is used, it is very difficult to extract the resolution enhancement from a priori considerations. In this paper we propose a protocol and experimental setup for the resolution measurement, demonstrating the resolution enhancement for different collection PSF values.

## 1. Introduction

The study of living cells or cell subcellular organelles requires a microscopy technique at the same time safe and providing high resolution. The fluorescence microscope is an excellent tool as it allows to identify cells and submicroscopic cell components with a high degree of specificity both in vitro and in vivo [1]. A variant of the fluorescence microscope is confocal fluorescence microscopy [2] [3], whose principle of operation is to prevent the detection of out-of-focus light by placing a pinhole aperture between the objective and the detector to let only the focused light rays pass [4]. The pinhole aperture must be much smaller than the Airy disk (the central region of blurry pattern generated on the image plane of an optical microscope by the signal) to increase the resolution, however with such a small aperture useful signal may be lost. [5]. To get around this limit, super-resolution techniques based on deconvolution are applied to the resulting images [6]. Deconvolution is mainly concerned with solving image degradation which is essentially due to the diffraction of light when it passes through the optical path of the imaging system [7]. The degraded image is the product of convolution between the real image and the PSF (Point Spread Function) of the objective [8], a function that models the three-dimensional diffraction generated by a point light source passing through the imaging system [9], thus the use of deconvolution is required to reverse the process and get the real, non-degraded image. SIM is a super-resolution technique that illuminates the sample with spatially structured illumination patterns, therefore known, so the image obtained is the product of the known model and the unknown structure of the sample. Through this technique it is possible to access high resolution information of the sample [10]. However, SIM is sensitive to the sample and to aberrations due to illumination, furthermore the experimental setup is extremely complex [11]. To solve these problems, we can use the blind-SIM technique, which reconstructs the image without knowing illumination patterns [11]. A variant of blind-SIM is the SAI technique (Scattering Assisted Imaging), which is capable to improve the imaging resolution by exploiting an illumination PSF (speckles grain size), smaller than the collection PSF (defined by the collection objectives) [12]. However, if SAI is used, it is very difficult to estimate the resolution in a specific experimental configuration a priori,, that is starting from the knowledge of the illumination PSF and the focusing PSF [13], hence in this paper we propose a protocol for the resolution measurement by adding an iris to vary the collection aperture without changing the speckles grain size to obtain a low-resolution image to be improved with SAI and subsequently measure the ratio between the low resolution and improved resolution.

## 2. Results

Here, the resolution measurement is obtained by running SAI technique on images of merged fluorescent beads seen under widefield fluorescence microscope. The sample seen under the microscope is a dilute solution of ThermoFisher Nile Red fluorescent beads of diameter 250nm. After putting 5 ml of solution on the slide, we place it on the hotplate for 1 min at 50°C in such a way that the particles at the edge of the drop merge to create elongated structures that intersect each other. In our study we will look at the intersection between two elongated structures to measure their resolution. To illuminate the sample with speckles we use a laser beam with a wavelength of 532 nm and a digital micromirror device (DMD) that modulates the laser beam to generate speckles which pass through the telescope consisting of the L1 lens and the objective. The beam, after passing through the L1 lens, reaches the objective through a dichroic mirror. The speckled laser beam excites the beads that emit fluorescent light with wavelengths in the range 540-750 nm. Since the laser used does not excite at a single wavelength due to its low quality, we put a 550 nm long pass filter before the CCD camera to be sure to collect only fluorescent light. The fluorescent light of the sample is passed through the dichroic mirror, reaches the mirror which deflects the beam in the direction of the camera to be viewed through the L2 lens. Between the mirror and the L2 lens we place an iris. We modify iris’ aperture to increase the size of the collection PSF (Fig.2b, c) and therefore decrease the resolution of the image, on which we run SAI to increase its resolution. Experimental setup is shown in Fig.1. Our goal is to see how much resolution improves using SAI as the ratio *η* changes

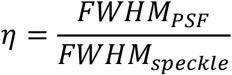

**Fig.1.**
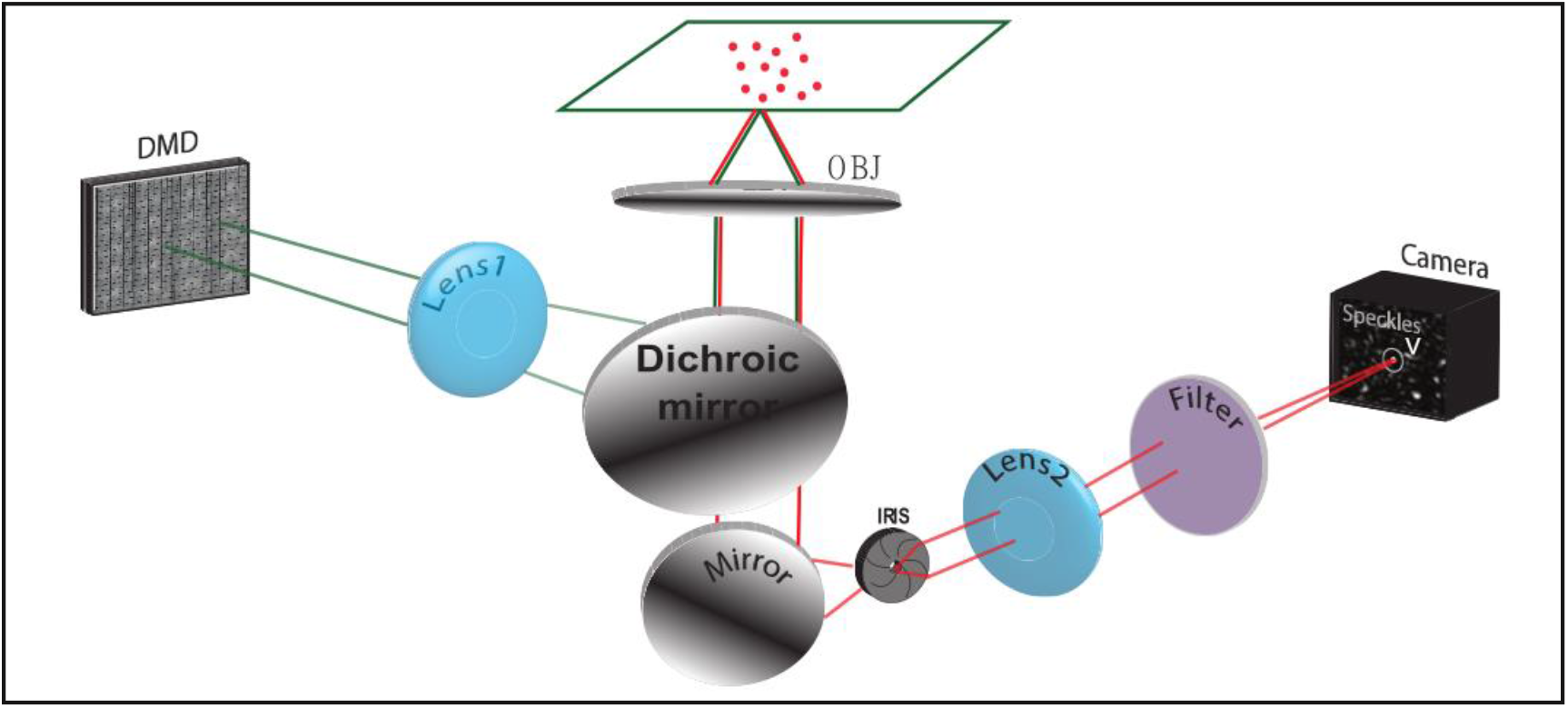
Experimental setup: The wavefront of a laser beam (λ = 532 nm, laser waist 0.7 mm, divergence < than 0.5 mrad) is modulated by a digital micromirror device (DMD, 1024 × 768 micromirrors, pixel size of 13 μm) to generate a speckled beam. The active area (50 × 50 micromirrors) of the DMD is imaged with a demagnification factor of 11 at the sample plane through a telescope composed by lens L1 (f = 200 mm) and the illumination/collection objective (OBJ). Fluorescent light from the sample is collected through a dichroic mirror (DH), passed through an iris, and imaged on a CCD camera through a lens L2.

The observable variable *η* is an indicator of the goodness of the collection compared to a fixed illumination (in our case the speckles). The FWHM of the speckle (Fig.2a) does not vary with the variation of the iris aperture as it depends only on the sample.

**Fig.2.**
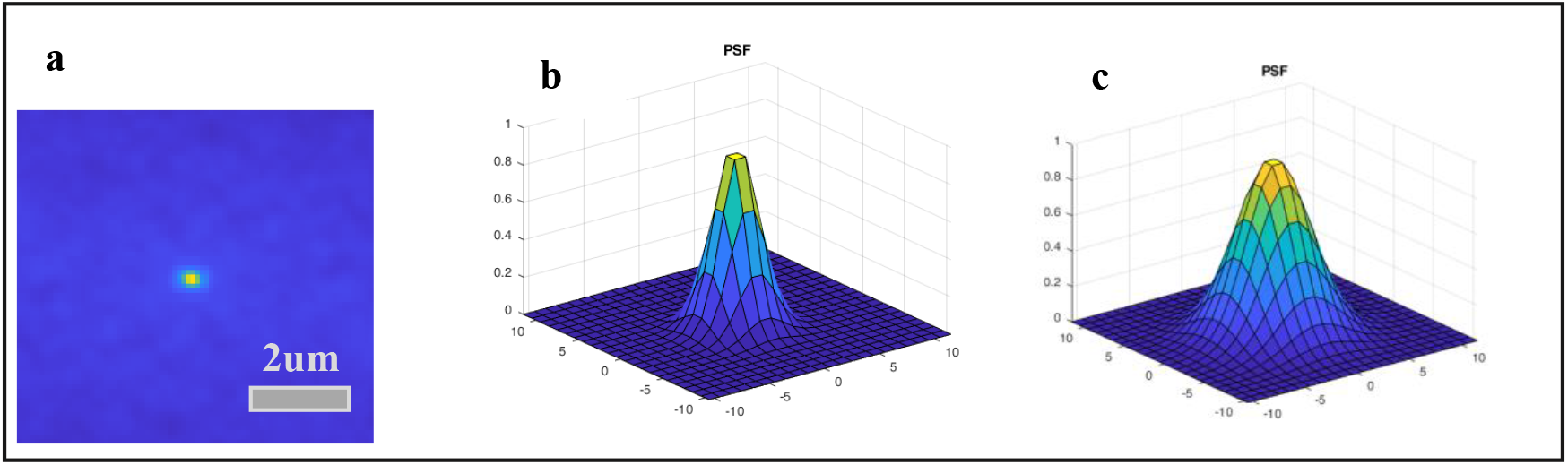
(a) Speckle (FWHM_speckle_= 3.1571). (b) PSF when iris is fully open (FWHM_PSFo_=4.0232). (c) PSF when iris is almost completely closed (FWHM_PSFc_=6.8634).

We run the standard SAI protocol: we acquire 600 frames when the iris is fully open, we make the average which will represent the high resolution reference image, the Ground Truth (GT). Then we acquire 600 frames when the iris is almost completely closed, and we make the average, and we get the low resolution image (LR). On the LR we run SAI and we will get a figure that we will call SAI’s IMAGE (Fig.3).

**Fig.3.**
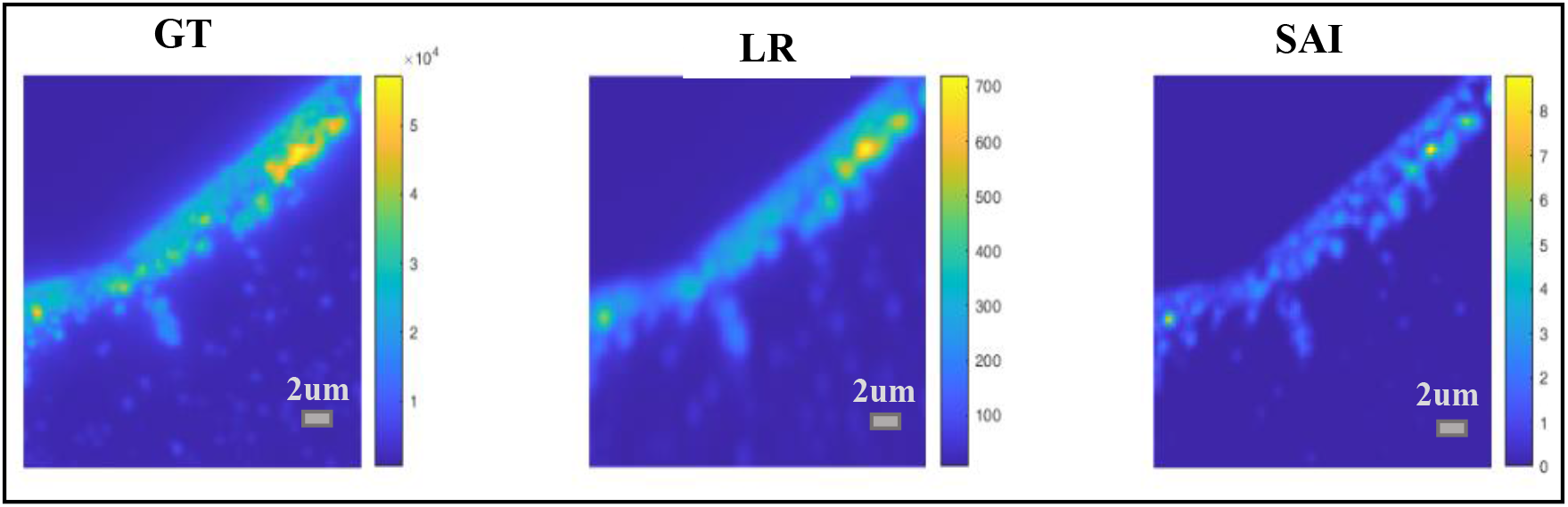
Comparison between GT, LR and SAI’s IMAGE.

We consider two elongated structures consisting of fused beads (fig.4a) to see where they are resolved in the case of the low resolution image (LR) and in the case of the image obtained with SAI (SAI’s IMAGE). We trace the intensity profile at twenty points along the axis that passes horizontally through the two elongated structures. For each intensity profile we evaluate the maximum intensity peaks with respect to the central dip and see in which points the two elongated structures are resolved according to the Rayleigh criterion (they are solved when the central dip is about 26% of the maximum intensity) [14] both in the low resolution case and in the SAI case. Finally, we trace on SAI’s IMAGE the two lines that represent the intensity profiles at the points where the two images (LR and SAI’s IMAGE) are resolved (fig.4b), the yellow line passes through the point where the two elongated structures are resolved in LR and the red line passes through the point where the two elongated structures are resolved in SAI’s IMAGE.

**Figure5.**
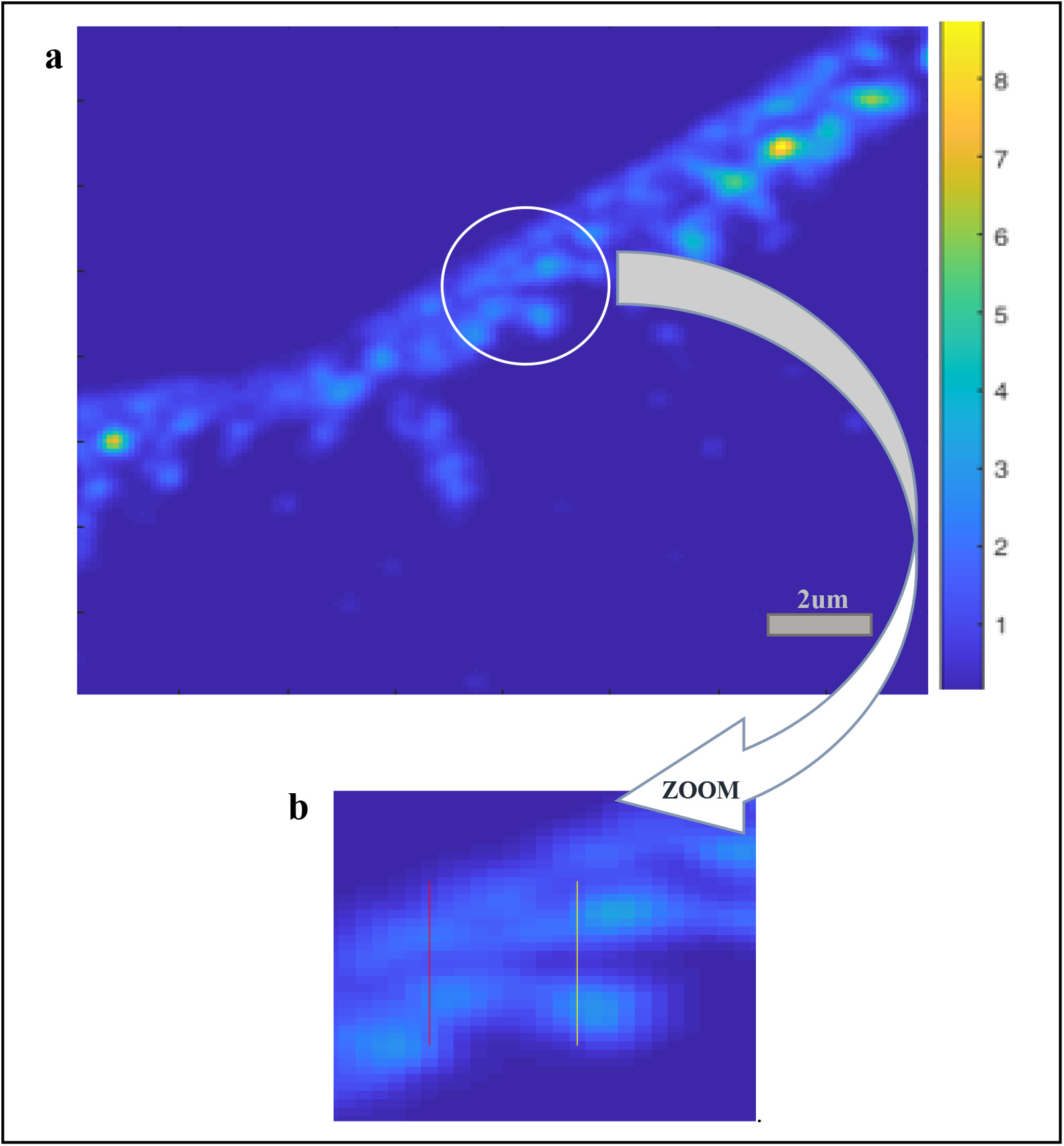
(a) we highlight the part in SAI’s IMAGE that we consider for our study and then zoom it (b) the yellow line passes through the point where the two elongated structures are resolved in LR and the red line passes through the point where the two elongated structures are resolved in SAI’s IMAGE.

In the case of LR, the resolution distance is 11 pixels (Fig.5a), or 1.7875 μm as the image pixel size is 0.1625 μm obtained from the ratio between the camera pixel size and the magnification of the lens. In the case of SAI’s IMAGE, the resolution distance is 7 pixels, or 1.1375 μm (Fig.5b).

**Figure5.**
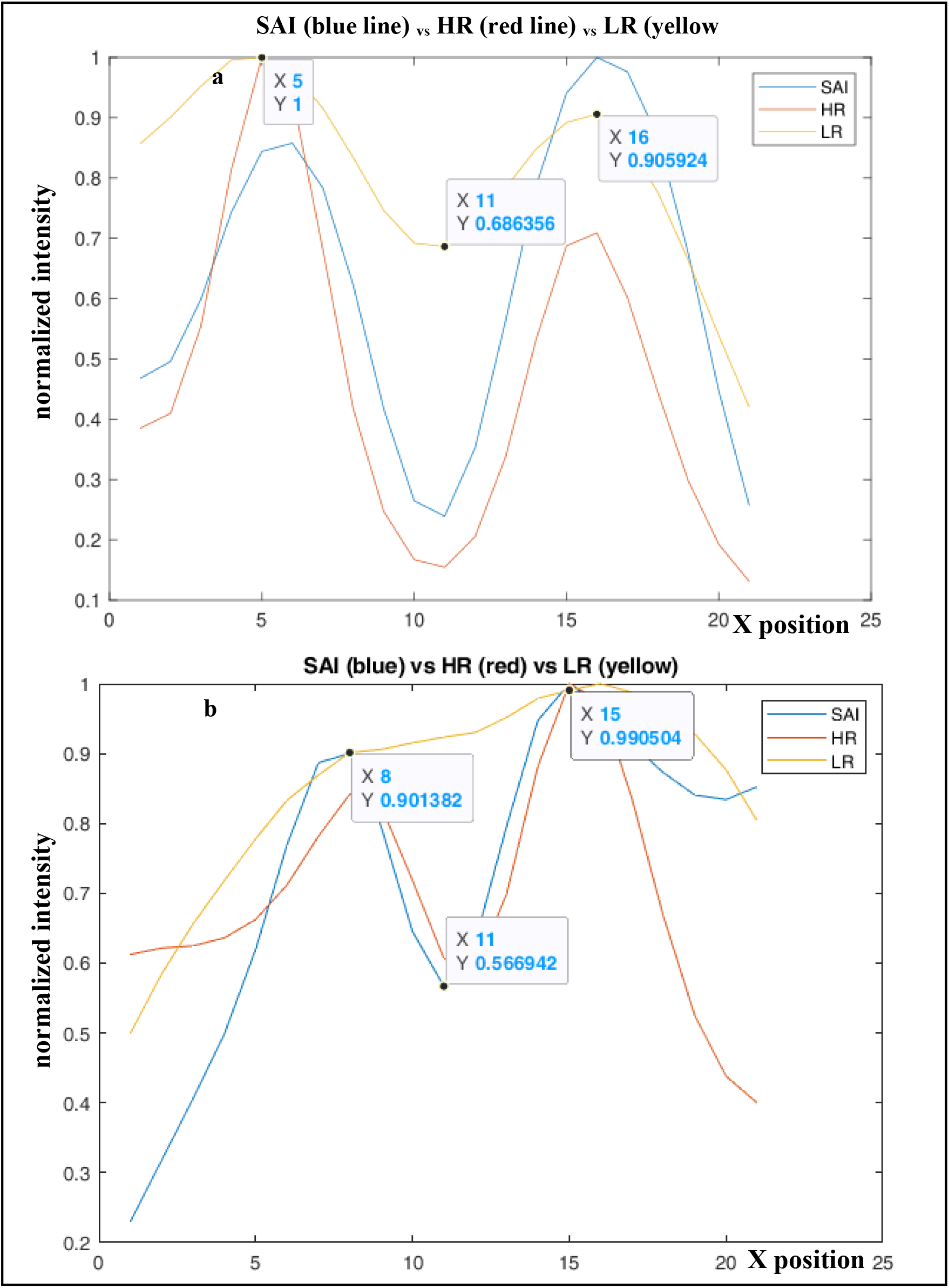
(a) Intensity profile when LR is resolved. (b) Intensity profile when SAI’s IMAGE is resolved.

The ratio between the two resolutions indicates how much resolution is increased using SAI.

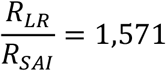

## 3. Conclusion

30 experimental tests were performed with 30 different samples by varying the dilution and the volume of the diluted solution on the slide, in 96% of cases we obtained the results shown in this study, in the remaining 4% or we had diluted the beads too little, so they emitted too much light and it was not possible to distinguish the intersected elongated samples, or the volume of the diluted solution was so high that it did not dry in the expected time and therefore the beads of the droplet outline did not melt. The effectiveness of the protocol and its simplicity, because for measuring resolution requires only the use of fluorescent beads and an iris with variable aperture, make it desirable. This technique is not sensitive to aberrations due to illumination but it is sensitive to those due to the collection therefore solving this type of aberrations with liquid lens [15], the effectiveness of this protocol will improve.

## Notes

### Competing Interest Statement

The authors have declared no competing interest.

### Summary of Updates

for adding an author: Emmanouil Xypakis

